# ML-guided robotic microinjection of single neurons in human brain organoids

**DOI:** 10.64898/2026.02.16.706073

**Authors:** Martina Polenghi, Jacob O’Brien, Elena Restelli, Suhasa B. Kodandaramaiah, Elena Taverna

**Affiliations:** Human Technopole, Viale Rita Levi Montalcini, Milan, IT; Department of Mechanical Engineering, University of Minnesota, Minneapolis, USA; Graduate program in Neuroscience, University of Minnesota, Minneapolis, USA; Department of Biomedical Engineering, University of Minnesota, Minneapolis, USA

## Abstract

Human brain organoids have emerged as powerful models for studying the physiology, pathology, and evolution of the human brain. When combined with single-cell approaches, they offer the potential to directly probe the dynamics and mechanisms underlying cell fate decisions. A major technical bottleneck, however, remains the ability to reliably visualize and manipulate individual cells within their dense and heterogeneous tissue environment. Microinjection has proven effective for this purpose, allowing direct delivery of membrane-impermeable probes into single cells within intact tissue. Despite its versatility, widespread adoption of microinjection has been limited by its technically demanding and low-throughput nature. Automated microinjection systems developed for murine tissue have demonstrated that robotics can overcome these limitations, enabling systematic single-cell lineage tracing at scale. Here, we present a vision-guided robotic system capable of imaging organoid slices, identifying tissue boundaries, and targeting specific single cells for microinjection. This approach is generalizable across murine and human tissues and enables high-throughput single-cell manipulation, opening new avenues for studying human brain development at scale.

## INTRODUCTION

Studying how single cells functionally contribute to the development of a whole tissue is a fundamental question in developmental biology. In developmental neuroscience, this question has been traditionally addressed *in vivo* using animal models or *in vitro* using organotypic brain tissue (Humpel 2015). Of note, the reconstruction of cellular history in human tissue is a challenging and often unattainable task, due to technical and ethical limitations. Recent advances in iPSC derived technologies provide a powerful way to study these processes in humans, as organoids can recapitulate development *in vitro* (Takahashi and Yamanaka 2006). Organoids are formed by cells that self-organize in 3D and provide an accessible way to study developmental patterns and trajectories in human tissue, reproducing (at least partially) its tremendous complexity (Sloan et al. 2018; Lancaster and Knoblich 2014; Qian et al. 2020). In the field of developmental neuroscience, the advent of human brain organoids has represented a major step towards the understanding of neurodevelopmental mechanisms across physiology, pathology and evolution. Combining human brain organoids with single cell manipulation techniques would provide a lens directed on the dynamics, mechanisms and molecular logic underlying cell’s fate choices and trajectories during brain development. A challenge in that respect is the ability to clearly visualize single cells from the intricate complexity of the surrounding tissue.

This has been successfully done via microinjection, a technique that involves the insertion of a fine-tip glass capillary filled with membrane-impermeable compound into single cells. In previous studies (Taverna et al. 2012; Wong et al. 2014) brain organotypic slices have been microinjected to manipulate and characterize single neural stem cells in intact tissue from mouse, ferret and human. Its flexibility makes microinjection amenable for high-resolution lineage tracing, gene editing and morphological analysis. However, large scale, systematic studies are precluded by the tedious and skilled nature of microinjection. Efforts have been made to automate this process in murine tissue, where automated microinjection has been developed (Shull et al. 2019; 2021) to overcome the limitations of manual microinjection, thereby increasing the success rate of this procedure and decreasing the differences due to user-dependent skills. This allowed to reconstruct the lineage of microinjected apical progenitors and to follow their fate and identity successive to cell division, acquiring a privileged picture of cellular dynamics occurring during mouse brain development.

Extending the use of microinjection automation to brain organoids would allow to directly study human brain development at the single cell level. Recently, progress has been made in utilizing robust but generalizable machine learning algorithms for object detection in vision guided robotic microinjection (Alegria et al. 2024; Joshi et al. 2021). Our work extends such approaches to perform single cell microinjection in intact tissue, targeting single cortical neurons in human iPSCs-derived brain organoids. We report on a ML vision guided robot that can automatically (i) image organoids brain slices (ii) detect tissue boundaries and interfaces and (iii) inject into single cell at specific locations within the organoids. We show that this approach can be generalized to both mouse samples and human iPSC derived brain organoids, where we successfully injected cells and reconstructed their morphology and intracellular architecture in tissue context. Notably, this can be done at scale, with the ability to microinject an average of 1.76 cells per second in a single experimental session, depending on the tissue edge’s length and on the targeted focal planes.

## RESULTS

### Human Brain organoid as model system

Human brain organoids represent the state-of-the-art model to recapitulate and study *in vitro* the cellular and molecular logic of human brain development in physiological and pathological context (**Fig. 1A-D**).

**Figure 1.**
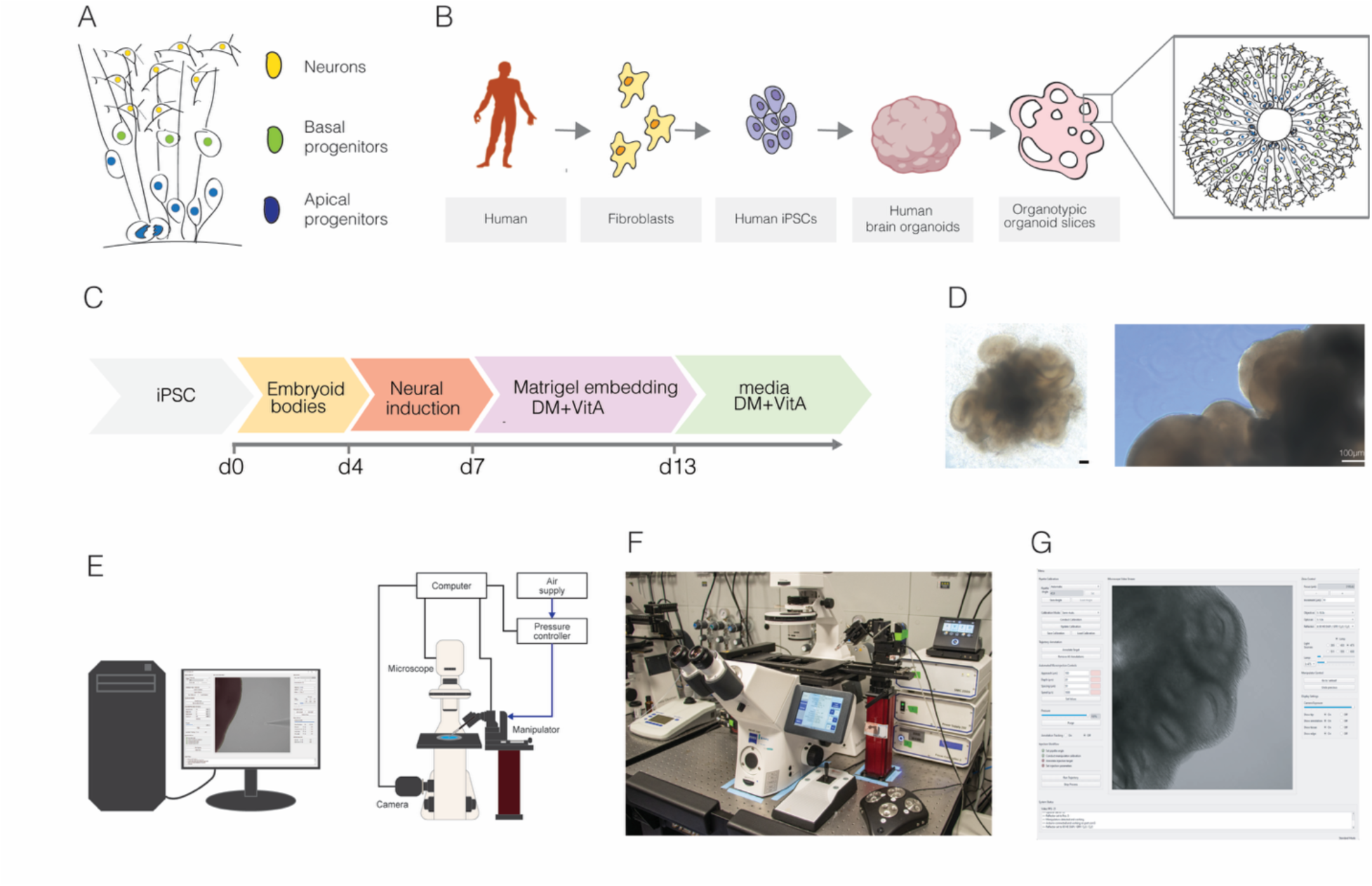
Automation to address human biology at single cell resolution. In mammals, neurons forming the brain are generated during embryonic development from neural stem and progenitor cells (A). The advent of iPSCs technology (B) allows for the modeling of human brain development in vitro in healthy and affected individuals using human brain organoids (C-D), that mimic and recapitulate key steps of the developmental process (Scale bar 100 µm). To study development at the single cell level, we developed a robotic tool (hardware and software) to target and manipulate the different cell populations present in the organoid. (E) Schematic representation of the automated microinjection system showing the core elements of the system (F) Photograph of the autoinjector system showing the physical layout. (G) Representative image of the graphic user interface of the ML-guided Autoinjector control software.

We adapted the protocol developed by Lancaster et al. (Lancaster and Knoblich 2014) to obtain brain organoids from WTC11 iPSC line (**Fig. 1C-D**). Human brain organoids at day 40 (d40) showed an area of highly organized and densely packed cells forming the ventricular zone (VZ) and a less compact region reminiscent of the sub-ventricular zone (SVZ) in the primary embryonic tissue (**Fig. 2A**). The identity and distribution of neural progenitors (NPCs) and neurons were assessed using fate markers. The VZ contains Sox2-positive apical progenitors (APs; aRGs; **Fig. 2B**) and the SVZ is mainly constituted by Tbr2-postive basal progenitors (BPs; IPs and bRG/oRGs; Figure 2B). We investigated which neuronal subtypes were present by staining for NeuN (a pan neuronal marker), Tbr1 (marker for layer VI and I neurons), Ctip2 (marker for layer V neurons) and Satb2 (markers for layer IV neurons). From a qualitative point of view, irrespective of the neuronal marker used, neurons appeared located basal to the VZ and to the SVZ area, suggesting a correct cytoarchitecture of the human brain organoid (**Fig. 2C-E**). The quantifications showed that at Day 47 organoids were populated by Tbr1- and Ctip2-positive neurons, while only a fraction of neurons was Satb2-positive (**Fig. 2F**). We quantified the proportion of neurons double positive for Ctip2 and Tbr1 or for Ctip2 and Satb2 (**Fig. 2G**) and found that neurons positive for both Ctip2 and Tbr1 were approximately 100% while approximately an 80% of Satb2 neurons were also positive for Ctip2 marker. These data show that human brain organoids contain layer VI, V and VI neurons and suggest that, though simplified in terms of organization and cellular composition, this model represents an amenable alternative to primary human tissue to study brain development.

**Figure 2.**
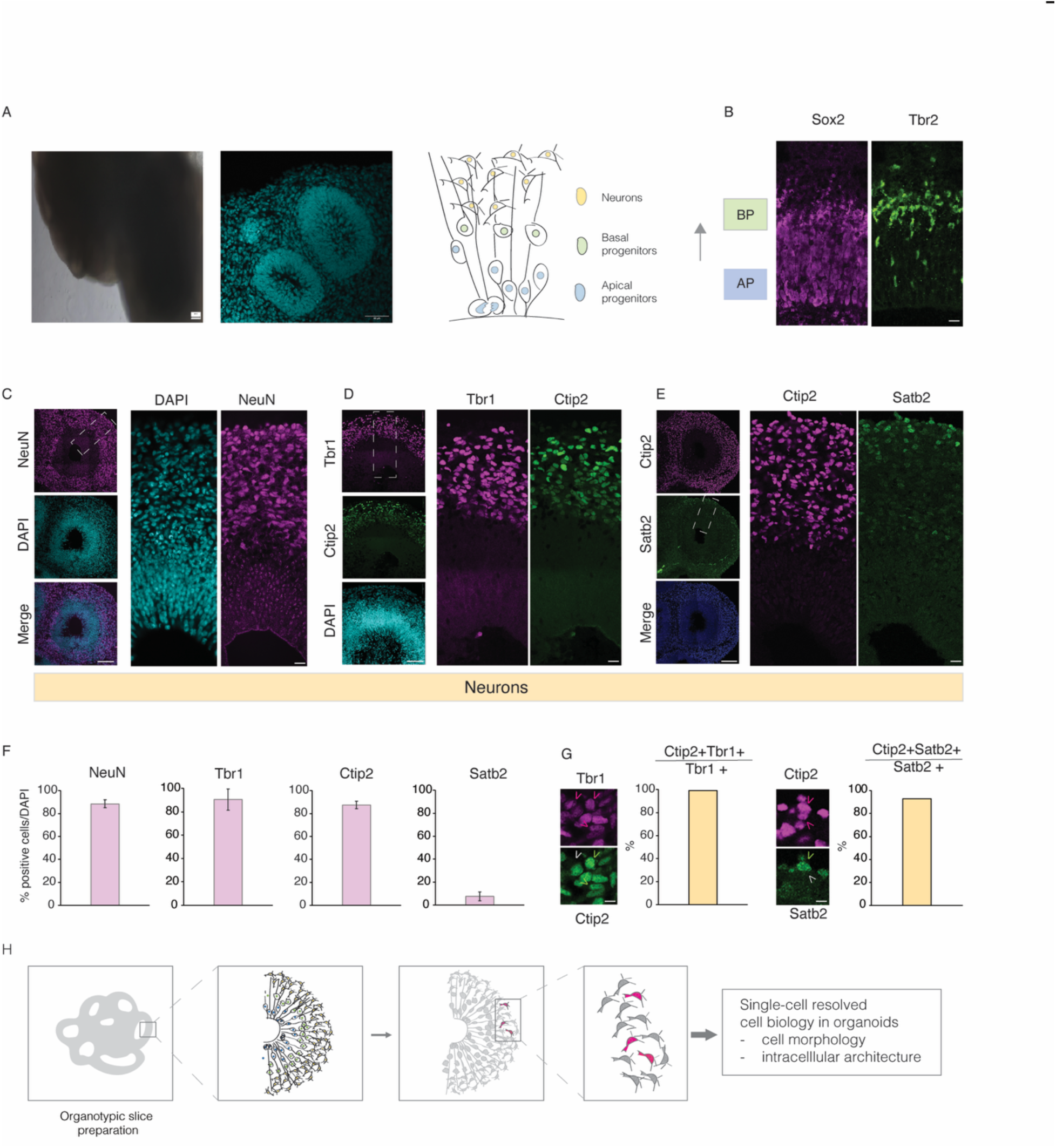
Human brain organoids model human brain development in a dish. Human brain organoids derived from iPSCs following the Lancaster protocol appear as sphere-like structures and contain a variable number of ventricles (A) Scale bar: 50 µm. (B) Apical and basal progenitors are found at different location and distance from the ventricular surface. Scale bar: 20µm. (C-E) APs and BPs give rise to NeUN and Tbr1 positive neurons expressing layer/specific markers such as CTIP2 and SATB2. Scale bars: 50-20 µm. The percentage of NeUN, Tbr1, CTIP1 and SATB2 positive cells is shown in (F) (expressed number positive cells/total DAPI). In (G), CTIP2 and SATB2 double cells were quantified. Scale bars: 10µm. The complexity of the human brain organoids calls for a technology to directly target neurons in single ventricles for the combined assessment of neuron’s identity, morphology and intracellular architecture (H).

### A robot for microinjecting single neurons in intact brain organoids

Crucial to the success of single cell studies are methods allowing to clearly visualize and separate single cells from the intricate complexity of the surrounding tissue. Being able to discriminate single cells in a crowded tissue is particularly critical for the study of single cells dynamics, as well as for reconstructing their cellular history and contribution to tissue morphogenesis. This is typically done via microinjection, a technique that involves the insertion of a fine-tip glass capillary of membrane-impermeable compound into single targeted cells. Microinjection has been successfully used to label or manipulate cells in 2D culture (Zhang and Yu 2008), and more recently, in organotypic slices from the developing brain (Taverna et al. 2012; Wong et al. 2014). However, microinjection is a low-throughput and low-yield technique that requires precision and practice for the user to master. Recently, we had demonstrated that robotics, equipped with computer vision algorithms could automatically target intact 2-dimensional brain tissue slices (Shull et al. 2019; 2021) as well as whole intact embryos (Alegria et al. 2024; Joshi et al. 2021; Guo et al. 2024). Here we adapted this robotic microinjection methodology for targeting neurons in intact iPSCs-derived human brain organoids slices to study cell morphology and intracellular architecture of single targeted cells.

The microinjection robot (**Fig. 1E-F**) consists of motorized XYZ pipette manipulator integrated into a standard inverted epi-fluorescence microscope. The microscope has an integrated XYZ stage that allows manipulation of the brain organoid tissue independent of the micropipette. A computer-controlled pressure regulator adapted from our previous work (Kodandaramaiah et al. 2018) is used to programmatically regulate the internal pressure of the pipette during microinjection.

The microscope camera images the tissue and the pipette. The robot utilizes computer vision algorithms to perform real-time detection of the tissue surface and the pipette tip. Once in focus, the tissue can be automatically annotated via a graphic user interface (GUI) that allows the user to input the desired parameters for targeted microinjection (**Fig. 1G**). The tip of the micropipette needs to be sequentially moved to target locations in tissue to execute the microinjection. Machine-lead movement of the pipette in 3D cartesian space results in displacement of the pipette in the microscope field of view (FOV). As the target locations are in the image, the robot automatically performs a calibration process that accurately translates the movement of the pipette in 3D cartesian space via the motorized XYZ stage, to motion in 2D pixel coordinates in camera FOV. The robot then manipulates the tip of the micropipette to the desired injection location at the tissue interface, briefly inserts the pipette to a specified depth inside the tissue while under controlled internal pressure and retracts the pipette to complete the microinjection process. The process is sequentially performed along the length of the tissue surface at each desired microinjection target location. The application of this robotic platform on human brain organoids allowed us to obtain single-cell resolved observations on targeted cells’ morphology and intracellular architecture (**Fig. 2H**).

### Real-time computer vision detection, focusing and calibration of microinjection pipette

To achieve full automation of single cell microinjection in intact tissue, we needed the robot to (a) automatically detect the location of the tip of the injection micropipette in the camera field of view (FOV) and bring it into focus, (b) move the pipette to a desired location within the FOV, (c) image tissue, and automatically identify target locations of interest for microinjection, and (d) have the ability to sequentially target multiple locations on the tissue accurately and microinject the cells (**Fig. 3A**).

**Figure 3.**
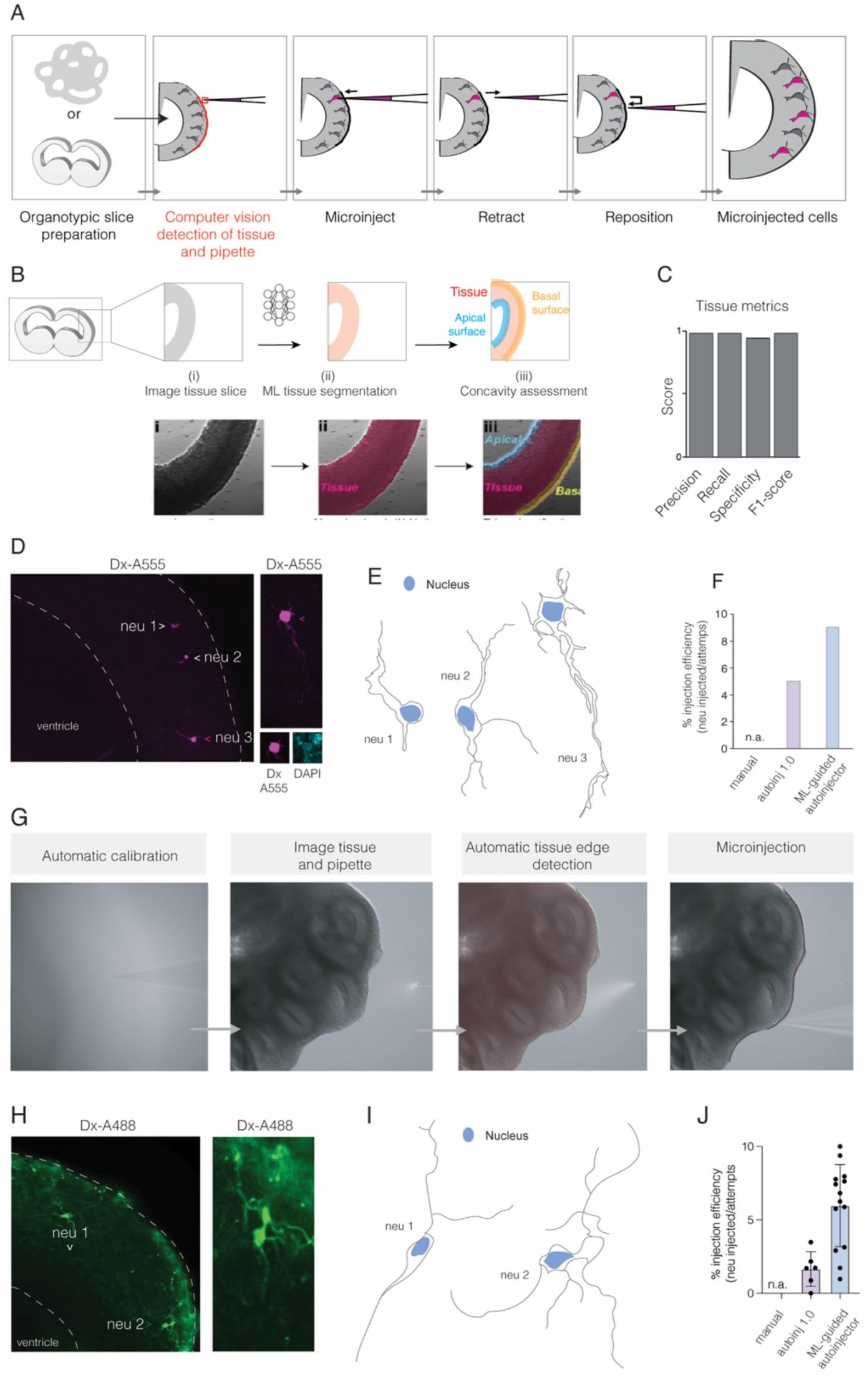
Autoinjector operation. (A) Schematic representation of the overall automated micro injection process. Images of tissue slices are annotated using computer vision to identify the locations of the tip of the auto micro injection needle and the tissue interface. Once this is determined, a trajectory that the micro injector needle tip follows is computed, and cells at a given anatomical location and tissue are sequentially targeted for microinjection. Breakdown of the computer vision detection of tissue is shown in (B). A machine learning model is used to segment the tissue, followed by a concavity assessment to identify the basal and apical surfaces. (C) Quantification of the tissue segmentation metrics. (D) Representative image showing neurons in organotypic murine brain slice automatically injected using the ML-guided autoinjector (neurons in magenta). Scale bars: 20 and 5 µm. (E) Morphology of single neurons was revealed by automated microinjection in the mouse telencephalon. (F) Comparison of microinjection efficiency of the current ML-guided autoinjector to the previous version of the auto-injector (Shull et al. 2021; 2019) that did not use machine learning-based segmentation. (G) Representative images during an automated microinjection experiment of human brain organoids. Images show sequentially – Calibration, tissue and tissue segmentation and edge detection and one microinjection attempt. A representative image of neurons automatically microinjected with the dextran A488 dye in a human brain organoids (neurons in green). Scale bars: 20 µm. (H) Morphology of single neurons was revealed by automated microinjection in human brain organoids. (I) Quantification of injection efficiency in human brain organelles using the current robot and the previous robot. No manual injections were successful.

We trained neural network (YOLOv5) to detect the X, Y, and Z positions of the pipette tip. The neural network output XY pixel position of the pipette tip in the image and a Z classification of whether the pipette is “below”, “in”, or “above” the current focal plane. The autoinjector then used the Z classification to iteratively raise/lower the pipette tip until it is in focus. We characterized the accuracy of the pipette tip detection algorithms by first evaluating the z-classification algorithm for detecting the focus plane compared to that determined by a human (**Supplementary Fig. 1A**).

Once the pipette was detected and in focus, it was automatically moved to a series of known locations in cartesian coordinate space, and the displacement of the tip of the pipette in pixel coordinates was automatically detected. A transformation matrix related the position of the pipette in cartesian coordinates to displacement in the FOV (**Supplementary Fig. 1B**). This calibration step allowed us to position and manipulate the micropipette within the microscope imaging field-of-view with an accuracy of <5 µm (**Supplementary Fig. 1C**).

### Real-time annotation of microinjection targets and tissue displacement tracking

Automatic target annotation was achieved using a two-step process. First, neural network tissue segmentation algorithm categorized the tissue versus the background (**Fig. 3B**) followed by an edge classification algorithm that evaluated the concavity of the detected edge to categorize it as apical or basal based on their curvature (**Fig. 3B**). We used a U-Net model (Navabi et al. 2025) to categorize every pixel in an input image as belonging to a piece of tissue or to the background. The U-Net returned a binary image where each pixel value corresponds to 0 (background) or 1 (tissue). This binary image was then used for edge classification model where detected tissue boundaries are extracted and analyzed for their concavity: concave edges (where the edge surrounds/encases background pixels) are classified as apical and convex edges (where the edge surrounds/encases tissue pixels) are classified as basal. Across all metrics, including precision, recall, specificity, and F1 score, our overall tissue segmentation scores were >0.9, indicating very high accuracy of detecting the requisite tissue surfaces for microinjection (**Fig. 3C**).

A final issue that confounds automated microinjection is tissue displacement and drift due to the mechanical interaction between the pipette and tissue slices. Tissue displacement is undesirable because it results in the injection target annotation at the beginning of the microinjection process to no longer being coincident with the tissue thereby resulting in failed injection attempts. We implemented a real-time tissue displacement-targeting algorithm that converts the injection target annotation into a series of tracking points in the image and iteratively updates the tracking point locations to re-coincide with the displaced tissue. First, the injection target annotation is initialized either by the user manually drawing the annotation or by the automatic annotation process detailed above (**Supplementary Fig. 2**). This annotation is then discretized to a series of uniformly spaced points that will be tracked in the following steps. These tracking points correspond to pixel locations in the camera’s image that are likely near the tissue edge (assuming the annotation is drawn on the tissue edge). Then, when a new image is available from the camera, optical flow is used to estimate the new location of each tracking point. In brief, optical flow works by assuming that the intensity/brightness of the points being tracked is constant (i.e. the tissue doesn’t get brighter/dimmer as it moves); the optical flow uses this assumption to evaluate a series of mathematical expressions to estimate the motion of objects between consecutive image frames and uses this motion estimation to update the tracking point locations.

### Robotic microinjection performance in mouse brain slices and human brain organoids

Single-cell labeling techniques are useful for the analysis of single cell morphology, intracellular architecture and dynamics. All techniques available to date label neurons via the labeling of neuronal progenitors, and the subsequent inheritance of the fluorescent reporter in the downstream neuronal progeny. In this optics we sought instead to apply the ML-guided Autoinjector to directly target single neurons for downstream analysis.

We first evaluated the robot performance by injecting embryonic mouse brain slices cut at 250µm of thickness. The mouse tissue edges in the FOV were recognized and classified either as basal or apical basing on the concave/convex structure of the tissue. To target neurons, we selected the basal edge (**Fig. 3D**) and we successfully managed to sparsely label neurons with Dextran-A555 dye and to thus reconstruct their morphology (**Fig. 3E**). We also evaluated the injection efficiency comparing our ML-guided autoinjector with manual microinjection and with the previous version of the robot (Shull et al. 2021; 2019) and we observed an almost two-fold increase in success rate of microinjection into neurons when using ML-guided Autoinjector compared with Autoinjector (**Fig. 3F**). Of note, microinjection into single neurons within a tissue is not feasible using manual microinjection, making automated microinjection the only feasible method for targeting this cell type in a tissue context.

We next leveraged the ML-guided autoinjector to microinject human cortical brain organoids slices to dissect neuronal morphology. We prepared 250-µm-thick organotypic slices from human brain organoids and carried out microinjection with Dextran A488 10kDa by approaching the slice from the basal side (outer contour) (**Fig. 3G-I**). Though organoid shape is less homogenous than mouse tissue, the software correctly segmented the edges of the organoid slice and recognized the outer edge as basal. As we did for mouse tissue, we could successfully reconstruct neuronal morphology within human brain organoids (**Fig. 3I**).

We finally evaluated the microinjection efficiency on neurons from human brain organoids by counting the productive neuronal injections over the total of injection events, comparing between the Autoinjector 1.0 and the current ML-guided Autoinjector and we observed also in this case an increase in injection efficiency with the ML-Autoinjector (**Fig. 3J**).

### Reconstructing intracellular architecture of single cells in whole human brain organoids

We reasoned that by combining microinjection with immunofluorescence for intracellular organelles we could reconstruct the intracellular architecture of single human neurons in the 3D space. To this end, we optimized and developed a protocol for conventional immunofluorescence on floating sections obtained from brain organoids, and we also developed an alternative protocol for whole-mount immunofluorescence on the entire organoid, based on ultra-fast tissue clearing (**Fig. 4A**, see also Materials and Methods). We focused our attention of the Golgi apparatus (GA) as this organelle is studied in relation to cell polarity and cell migration. In apical progenitor cells (APs), a class of neural progenitors, GA is specifically positioned in the apical process of these cells, while being absent from the basal process. Upon cell fate switch and differentiation of APs into basal progenitors, the Golgi gets repositioned close to the nucleus. It has been observed that GA in mature neurons is polarized, being present in the cell body but absent from dendrites except for small Golgi outposts (Kemal et al. 2022; Ori-McKenney et al. 2012). To validate our microinjection experimental pipeline, we checked GA localization in microinjected neurons of human brain organoids, and we found that the GA is perinuclear, and it is absent from other cellular compartments (**Fig. 4B**). The GA mass is localized in the main process extending from the cell body, most likely the leading process of the migrating neuron.

**Figure 4.**
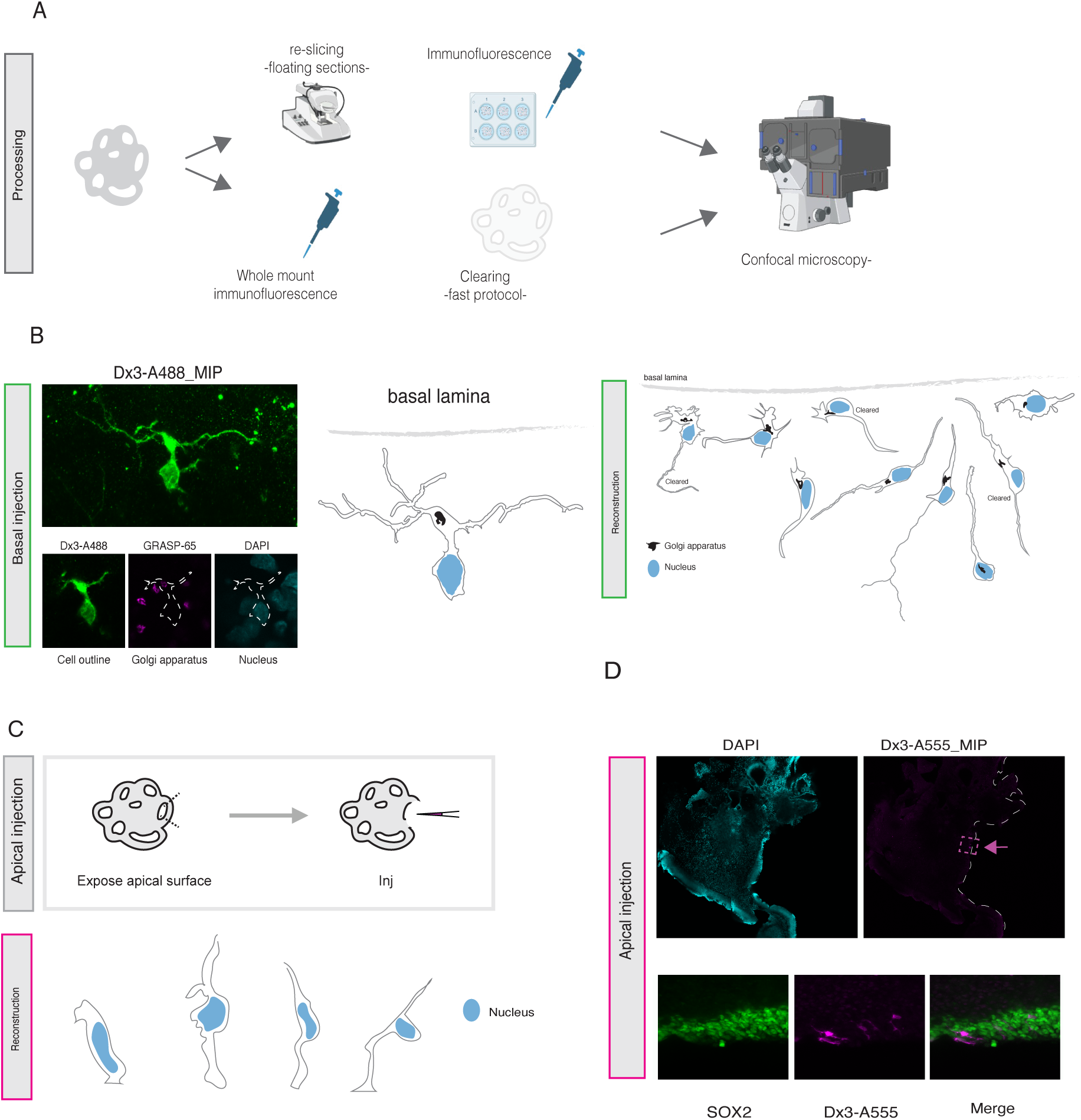
ML-guided automated microinjection in human brain organoids allows to reconstruct morphology and intracellular features in different cell types. (A) Schematic pipeline of organoids slices post-microinjection processing. We also developed an alternative protocol to stain whole slices from human brain organoids with a fast procedure for tissue clearing, without the need of re-slicing. This protocol allows for the full reconstruction of the 3D morphology and intracellular architecture of single microinjected human neurons and human neural progenitors. (B) Basal microinjection of neurons (green) in human brain organoids and co-staining for Golgi apparatus (GRASP65, magenta) allows to reconstruct cell morphology and Golgi apparatus localization within single neuronal cells. Scale bars: 10 µm (C) By exposing the apical surface of ventricles of human brain organoids, the ML-guided autoinjector can target apical progenitor cells and allow to reconstruct their morphology. (D) ML-guided microinjection of human brain organoids targeting single apical progenitor cells using Dextran A555 (magenta) and co-staining for SOX2 (green) as a marker for apical progenitors. Scale bars 100 µm and 10 µm.

Finally, we generalized the use of the ML-guided autoinjector to target human apical progenitors (APs) within the ventricular zone (VZ) of human brain organoids. We first dissected the organoid to expose the VZ and we microinjected Dextran-A555 along the surface of the exposed ventricle (**Fig. 4C-D**). Also in this case it was possible to target single sparse APs (double positive for dextran and Sox2, **Fig. 4D**) and to reconstruct their morphology (**Fig. 4C**, right part of the panel). The possibility to target apical progenitors allows to study in detail not only APs morphology and dynamics but also to track lineage progression during brain development.

In conclusion, ML-guided automated microinjection allows to gain insights into single cells morphology and intracellular architecture, providing a tool to study the cell biology and cellular dynamics of human progenitors and neurons in brain organoids.

## DISCUSSION

### A machine learning-guided microinjection robot to target and study single cells in the developing brain

Single-cell analysis techniques are at the forefront of neurodevelopmental studies as they allow to reconstruct single cell trajectories and cellular history. This is extremely relevant to define the cellular logic underlying tissue morphogenesis in contexts such as brain development where the timing of appearance of neural progenitors and neurons has a great impact on the overall neocortex formation. The use of human brain organoids provides researchers with the opportunity to study with unprecedented detail human brain development at the single cell level, as shown by seminal work employing scRNAseq to describe cellular trajectories in health and disease.

Some limitations underlie the techniques available to date, such as the fact that during tissue preparation and dissociation for e.g. scRNAseq, the cell is completely taken out of its tissue context, loosing not only connections with neighbours, but also its own morphology and overall architecture, features that are crucial in determining its proliferation/differentiation potential (Kalebic et al. 2019). Spatial transcriptomics techniques have significantly contributed to elucidate molecular regulatory mechanisms and cellular interplay within a tissue microenvironment, by preserving the tissue intact (Ke et al. 2025). Despite its increasing potential and applicability, spatial transcriptomics provides snapshots rather than information on dynamics, which are usually inferred computationally, thereby limiting its use when modelling lineage trajectories at single cell levels. Furthermore, spatial transcriptomics remains largely observational, precluding direct functional interrogation of individual cells in situ and limiting causal assessment of cell-cell interactions and fate decisions. This gap can be potentially addressed using microinjection, a technique that allows for precise targeting and manipulation of single cells. Although highly informative for the dissection of single cell contribution to brain development (Taverna et al. 2012; Florio et al., n.d.; Shull et al. 2019; 2021; Appiah et al. 2023; Ossola et al. 2024), microinjection present technical challenges, especially in heterogenous and complex tissues such as organoids.

The development of the robotic microinjection technique was prompted by the need to circumvent this difficulty in manipulating intact tissue at the single cell level. A key enabling feature of the approach presented here is the generalized ability to target tissue at specific locations using ML-guided object detection. While early efforts to use computer vision for automating single cell interfacing relied on sequential application of multiple image processing and recognition algorithms (Wu et al. 2016; Suk et al. 2017; Annecchino et al. 2017), recently machine learning algorithms have been generalized and used for object detection in vision-guided microinjection (Joshi et al. 2021; Alegria et al. 2024). Here we leverage these advances and developed a machine learning-guided (ML-guided) Autoinjector to automatize the entire pipeline of this experimental procedure, from pipette calibration to tissue edge detection and finally to serial microinjection into single cells. We trained our tissue detection model on mouse tissue slices and then extended the application on human brain organoids. Automatic annotation was achieved through a first neural network-based segmentation that allowed to distinguish the tissue from the background and then the edges were classified as apical or basal considering their curvature (concave vs convex). This process allowed to correctly detect the apical part and the basal side of mouse brain tissue and be later extended to human brain organoids. Of note, although human brain organoids are more heterogeneous and present fewer neat edges compared to the mouse tissue, our system did not need any re-training and could still recognize correctly the basal or apical surfaces, highlighting its robustness and generalizability. Based on these data, we predict that our model could be applied to other systems (both primary tissue and organoids) without significant time-consuming investment in re-training. The development of a real-time tissue displacement-targeting algorithm allows for on-line correction of tissue movement, making the system amenable for correcting tissue drift. This is particularly relevant for the targeting of primary tissues, organoids or even organisms that are spontaneously moving in space or contracting, such as cardioids or muscle tissue.

We applied our ML-guided Autoinjector to microinject single neurons in human brain organoids and we also tested its ability to target single apical progenitor cells from the apical side of the organoid. The injection efficiency (neurons injected over the total attempts) revealed an increase with the ML-guided Autoinjector compared to the previous version of the robotic platform, both in mouse and human tissue. Furthermore, we could successfully reconstruct the morphology and intracellular architecture of all microinjected cells in their native environment.

Overall, we believe that the ML-guided Autoinjector could be extended in future to single cell manipulation via injection of compounds to selectively impact cell behaviour, allowing to extend current methods for single cell tracking and manipulation. This would be particularly relevant to gain insights into cell biology and cellular dynamics of neural progenitors and neurons in different tissue types, among which human brain organoids to tackle cell biological questions in the context of development. Of note, the robotic platform we developed could be used beyond brain studies, to inject and functionally manipulate organoid types with complex architecture and heterogeneous cell composition, such as 3D cardiac organoids and liver models, allowing to dissect the role and contribution of specific cell populations to organ development.

## MATERIALS AND METHODS

### Mouse tissue collection

All animal studies were conducted in accordance with Italian animal welfare legislation, and the necessary licenses were obtained after approval by the Ministry of Health (project code 75DA4.N.B7F). Animals for these studies arrived from Charles River and were maintained in standardized hygienic conditions at Cogentech animal house in Milan. All experiments were performed on C57Vl/6J mice. Mouse brains were harvested at E14.5. embryos were transferred into phosphate-buffered saline (PBS). Embryos were decapitated using surgical forceps under a dissectoscope, skulls were open and brains extracted under visual control.

### iPSCs culture

The WTC11 iPSCs cell line was used to generate human brain organoids following the Lancaster protocol.

### Thawing iPSCs

To thaw iPSCs, the cryogenic vial was quickly put in the waterbath at 37°C for a maximum of 2 minutes. Cells were transferred, with a wet pipet tip, into a falcon already containing 5ml of mTeSR1 (Stemcell). The cell suspension was centrifuged for 5 minutes at 200xg. After centrifugation, the supernatant was discarded, and the cell pellet was resuspended into 1ml of mTeSR1 with RI. Then, 1/3 and 2/3 of cells were plated into new wells with a final volume of 2ml mTeSR1 with RI.

### iPSCs maintenance

iPSCs were cultured on 6-well plates coated with iMatrix and kept in mTeSR1 at 37°C and 5% CO2, the culture medium was changed daily. To maintain the iPSCs culture cells were passaged once a week, when about 80% confluency was reached. To passage cells, one wash with Dulbecco’s phosphate-buffered saline (DPBS) (Thermofisher), without calcium and magnesium (DPBS -/-) was done before incubation with 1ml ReLeSR (Stemcell) for 30 seconds; ReLeSR was then removed and cells were put at 37°C for 5 minutes. A 1ml of fresh mTeSR1 was added and used to detach cells through three to five washings to maintain intact colonies. Then the appropriate cell suspension was transferred to a new well, in a final volume of 2ml of mTeSR1 with iMatrix to allow cell attachment.

### Freezing iPSCs

To freeze iPSCs, the cells were washed with DPBS -/-, and incubated for 5 minutes at 37°C with 1ml of ReLeSR. Cells were then detached from the plate with 2 ml of mTeSR1 pipetting them for a maximum four to six times. The suspension was transferred to a falcon and centrifuged for 5 minutes at 200xg. The supernatant was discarded, and the cell pellet was resuspended with 500µl of mFreSR (StemCell), transferred to a cryogenic vial and progressively frozen to -80°C in a freezing box.

### Human brain organoid culture

Human brain organoids were generated following Lancaster et al. protocol (2015) with some modifications. Briefly, when WTC11 iPSC were at 70-80% of confluency cells were detached with TrypLE (Thermofisher) for 5 minutes at 37°C; 2 mL of mTeSR1 were then added to detach and resuspend in single cells. Cells were centrifuged for 5 minutes at 200xg and after discarding the supernatant they were resuspended in 1mL of mTeSR1 supplemented with RI. Cells were then counted and 9000 cells per well of a U-bottom 96-well plate were added to mTeSR1 with RI medium. Cells were then plated adding 200µL of suspension to each well of the 96-well plate to form Embryoid Bodies (EBs). Media change was done every other day using mTeSR1 without RI. At day 4, medium was changed to Neural Induction Medium (NIM DM-VitA was prepared with DMEM F12 (Gibco) in ratio 1:1, N2 1:200 (Thermofisher), 1:100 Glutamax (Thermofisher), 1µg/mL of heparin, 0.5% of MEM NEAA (Sigma) and 1:100 P/S) with a volume of 150µL per well. Media change was done every other day. At day 7 the EBs were embedded in Geltrex droplets and put in 6cm petri dishes in Differentiation medium without Vitamin A (DM-VitA was prepared with DMEM F12 and Neurobasal (Gibco) in ratio 1:1, B27 -VitA 1:100 (Thermofisher), N2 1:200, 1:100 Glutamax, 2.5µg/mL insulin (Sigma), 50µM 2-mercaptoethanol (Merck) and 0.5% of MEM NEAA and 1:100 P/S). Media change was done every other day until day 13 when the embedded EBs were transferred in DM+VitA (DM-VitA was prepared with DMEM F12 and Neurobasal in ratio 1:1, B27 +VitA 1:100, N2 1:200, 1:100 Glutamax, 2.5µg/mL insulin, 50µM 2-mercaptoethanol and 0.5% of MEM NEAA and 1:100 P/S). Organoids were then kept in culture until the desired age in shaking at 70rpm at 37°C.

### Brain organoid and mouse tissue slicing for single cell microinjection

Brain organoids or E14.5 embryonic mouse brains were embedded in low melt agarose 3% and cut at the vibratome with the tray filled with Tyrode (Sigma), with a thickness of 250µm. Organoid slices were collected in a 6cm petri dish in SCM medium and kept in the incubator at 37°C until use.

### Microinjection into human neurons in human brain organoids

For all experiments, microinjection was performed in pre-warmed CO2-independent microinjection medium (CIMM: DMEM-F12 (Sigma, Germany, D2906), 2 mM L-glutamine, 1X P/S, 1X N2 and B27 supplements, 25 mM (final concentration), Hepes-NaOH pH 7.3) (Taverna et al., 2012). Briefly, 1.5 µl of injection solution (5 µg/µl Dextran-Alexa488 or Dextran-Alexa555) was loaded into a microcapillary needle for microinjection. The microinjection needle was mounted on the capillary holder and microinjection was performed with the ML-guided Autoinjector by approaching the basal surface of human brain organoid’s slices.

Microinjected slices were transferred to a dish containing fresh slice culture medium (SCM) and transferred at 37°C in a humidified atmosphere of 40% O2 / 5% CO2 / 55% N2 for 30minutes before fixation.

The composition of the SCM is as follows: Neurobasal medium (Thermo Fisher Scientific, Germany), 10% rat serum (Charles River, Japan), 2 mM L-glutamine (Thermo Fisher Scientific), P/S (Thermo Fisher Scientific), N2 supplement (Thermo Fisher Scientific), B27 supplement (Thermo Fisher Scientific), 10 mM Hepes-NaOH pH 7.3.

### Fixation, embedding, and sectioning of microinjected organoids for immunofluorescence

Organoid and mouse tissue slices were washed twice with PBS to remove culture media, followed by fixation with 4% paraformaldehyde in 120 mM sodium phosphate buffer pH 7.4 at room temperature for 40 min and then stored in 1X PBS at 4°C overnight. Coronal sections of mouse slices (70 µm) were prepared from microinjected slices on a vibratome (Leica VT1000S).

All slices were collected in PBS and stored at 4°C for subsequent immunofluorescence staining.

### Brain organoid clearing for whole mount analysis

For the slices that were cleared we followed a previously published protocol (Shan et al. 2022). Briefly, for staining of 250µm thick brain organoid slices 30 minutes permeabilization with 0.5% Triton X-100 was done, followed by 30 minutes of blocking in IF buffer at room temperature. After blocking slices were incubated with primary antibody diluted in IF buffer at RT for 24h. The next day secondary antibody diluted in IF buffer were added to the slices for 24h at 80rpm. After washing antibodies slices were incubated with AKS (20% DMSO, 40% TDE, 20% sorbitol, and 6% Tris base dissolved in ddH2O and stirred at RT) for tissue clearing. The clearing solution was kept for 5 minutes at room temperature with shaking of 80rpm. Imaging was done right after clearing.

### Immunofluorescence

Floating sections were processed for immunofluorescence (IF) staining as previously described (Taverna et al., 2012). In brief, sections were permeabilized with 0.3% Triton-X100 (Carl Roth, Germany) in PBS for 30 min, followed by blocking in IF buffer for 1h at RT. The sections were then incubated with primary antibodies diluted in IF buffer overnight at 4°C. Subsequently, sections were washed 5 times for 5 min each time with IF buffer. Secondary antibodies were diluted in IF buffer with DAPI (Carl Roth, Germany) as nuclear counterstain. Sections were incubated at room temperature for 1h followed by washing (5 times 5 min each) with PBS. Stained sections were mounted in Mowiol (Sigma, Germany) and imaged. A complete list of all primary and secondary antibodies used together with corresponding concentrations are listed in table 1.

**Table 1.**

| Antibodies |  |  |
| --- | --- | --- |
| Rabbit anti GRASP65 polyclonal antibody (1:200) | Thermofisher | PA3-910 |
| Rat anti Eomes monoclonal antibody (1:200) | Thermofisher | 14-4875-80 |
| Goat anti SOX2 polyclonal antibody (1:200) | BioTechne | AF2018 |
| Mouse anti Dextran monoclonal antibody (1:250) | Stemcell | 60026 |
| Rabbit anti TBR2 polyclonal antibody (1:200) | Abcam | ab261913 |
| Rat anti Ctip2 polyclonal antibody (1:400) | Abcam | ab18465 |
| Rabbit anti Satb2 polyclonal antibody (1:200) | Abcam | ab34735 |
| Rabbit anti NeuN polyclonal antibody (1:200) | Invitrogen™/Thermofisher | PA5-78499 |
| Donkey anti-Goat IgG (H+L) Highly Cross-Adsorbed Secondary Antibody, Alexa Fluor™ Plus 488 | Invitrogen™/Thermofisher | A32814 |
| Donkey anti-Goat IgG (H+L) Highly Cross-Adsorbed Secondary Antibody, Alexa Fluor™ Plus 555 | Invitrogen™/Thermofisher | A32816 |
| Donkey anti-Goat IgG (H+L) Highly Cross-Adsorbed Secondary Antibody, Alexa Fluor™ Plus 647 | Invitrogen™/Thermofisher | A32849 |
| Donkey anti-Rabbit IgG (H+L) Highly Cross-Adsorbed Secondary Antibody, Alexa Fluor™ Plus 647 | Invitrogen™/Thermofisher | A32795 |
| Donkey anti-Rabbit IgG (H+L) Highly Cross-Adsorbed Secondary Antibody, Alexa Fluor™ Plus 555 | Invitrogen™/Thermofisher | A32794 |
| Donkey anti-Rabbit IgG (H+L) Highly Cross-Adsorbed Secondary Antibody, Alexa Fluor™ Plus 488 | Invitrogen™/Thermofisher | A32790 |
| Donkey anti-Mouse IgG (H+L) Highly Cross-Adsorbed Secondary Antibody, Alexa Fluor™ Plus 647 | Invitrogen™/Thermofisher | A32787 |
| Donkey anti-Mouse IgG (H+L) Highly Cross-Adsorbed Secondary Antibody, Alexa Fluor™ Plus 555 | Invitrogen™/Thermofisher | A32773 |
| Donkey anti-Mouse IgG (H+L) Highly Cross-Adsorbed Secondary Antibody, Alexa Fluor™ Plus 488 | Invitrogen™/Thermofisher | A32766 |
| Anti-Rat IgG(H+L) Secondary Antibody | Tissue Processing Facility Human Technopole |  |

### Confocal microscopy

Imaging was performed with confocal (Zeiss LSM 780 NLO; Zeiss, Germany) or widefield fluorescence (Zeiss Axio-something; Zeiss, Germany) microscopes. Images shown are 0.5 µm-thick single optical sections, unless stated otherwise. Confocal images were taken at either 63X or 40X magnifications. Immunofluorescence images of single optical sections were quantified using Fiji (Schindelin et al., 2012). Final image panels were processed with Photoshop and Figures were assembled using Adobe Illustrator CS5.1 (29.7).

### Image analysis and reconstruction of single cell intracellular architecture

We used a combined panel of cell morphological parameters and marker expression to score labelled cells in mouse tissue and in human brain organoids. Microinjected cells were identified as cells positive for the microinjection dye upon inspection at the epifluorescence microscope. To assess morphological organization and subcellular architecture, all positive cells in an experiment were imaged at high-resolution using confocal microscopy (see also above). The Golgi apparatus is detected using an antibody against the Golgi reassembly-stacking protein GRASP-65.

### Automatic annotation

The dataset for neural network tissue segmentation consisted of images of pipette tip randomly located at different XYZ locations in the microscope FOV. These images were captured with all combinations of the microscope optovar and objectives. These images were split 80% into the training data set and 20% into the testing dataset. The images in the testing dataset are not exposed to the network during training, so good performance on test images indicates the network may generalize well to novel data. All images were automatically annotated with the pipette tip XYZ position to serve as the “ground truths” for training.

Pipette tip detection error was evaluated by comparing the neural network pipette tip detection results to the annotated “ground truths” in the testing dataset. The neural network was first trained on the images in the training dataset. After training was completed, the trained network was used to detect the pipette tip position in the testing dataset. This detected pipette tip position was compared to the “ground truth” position to compute the positioning error.

The post-calibration positioning error was evaluated by first completing the calibration process (either automatically or manually). Then the computed calibration was used to send the pipette tip position to a random desired position in the microscope FOV. Then the user marked the current pipette tip position by clicking on the tip in the video feed to designate the actual pipette position. This actual position is compared with the desired position to compute the positioning error.

The calibration duration was evaluated by performing identical injection experiments on Autoinjector 1.0 and ML-guided Autoinjector and comparing the durations of each process. The injection experiment consisted of injecting a piece of tissue at three different focal planes and repeating this process for three pieces of tissue. The entire experiment (three focal plane injections for three tissue pieces) was timed, and the durations of each process were tabulated.

### Target annotation

The dataset for neural network tissue segmentation consisted of 40 pieces of embryonic mouse brain tissue that was imaged at three different focal heights for a total 120 images for neural network training. These 120 images were split 80% into the training data set (N=96) and 20% into the testing dataset (N=24). The images in the testing dataset are not exposed to the network during training, so good performance on test images indicates the network may generalize well to novel data. All images were manually annotated by a user to identify ground truth pixel values, so all pixels were manually labeled as “background” or “tissue”.

Tissue segmentation and edge classification performance were evaluated by comparing the neural network tissue segmentation and the edge classification results to manually annotated “ground truths” in the testing dataset. The neural network was first trained on the images in the training dataset. After training completed, the trained network was used to segment images in the testing dataset. The results of segmentation on testing dataset were used to count the number pixels corresponding to true positives (TP), true negatives (TN), false positives (FP), and false negatives (FN) in each image. These TP, TN, FP, and FN pixel counts were used to compute the tissue segmentation metrics.

The binary images resulting from tissue segmentation on the testing dataset were then input into the edge classification algorithm. An edge was classified as TP if it had the correct label (apical vs basal) and it coincided with the actual tissue edge. Edges were FP if they had an incorrect label or if it coincided with the background. Edges were FN if tissue segmentation/edge classification failed identify the presence of a tissue edge in the image. (TN were not counted because there are “infinitely” many non-existent edges in an image).

The annotation duration was evaluated by performing identical injection experiments on Autoinjector 1.0 and ML-guided Autoinjector and comparing the durations of each process. The injection experiment consisted of injecting a piece of tissue at three different focal planes and repeating this process for three pieces of tissue. The entire experiment (three focal plane injections for three tissue pieces) was timed and the durations of each process were tabulated and compared between Autoinjector 1.0 and ML-guided Autoinjector. Manual annotation durations (N=9) were tabulated from the Autoinjector 1.0 experiment and automatic annotation durations (N=9) were tabulated from the ML-guided Autoinjector experiment.

### Tissue tracking

Microbead tracking accuracy was evaluated by offline tracking the displacement of fluorescent microbeads on a coverslip as they were traversed through the FOV via the microscope’s motorized stage. First, the user smeared fluorescent microbeads on a coverslip, inserted the coverslip in the microscope, and focused the objective on the fluorescent microbeads. The user then took a video of the microbeads as they gradually translated the microscope stage. Microscope stage translation is gradual because the tissue is not dramatically displaced during the injection trial; typically, the tissue experiences a maximal displacement equivalent to the user-defined injection depth parameter which specifies how far the pipette penetrates into the tissue. The microscope system saves microscope metadata with each image frame in the video, so the microscope stage XYZ location is saved with each image. This metadata is used as the “ground truth” microbead displacement. After the video is acquired, the microbead displacements were analyzed with the tracker: first the user specified tracking points corresponding to the microbeads in the initial image, and then user ran the tracking algorithm on each subsequent image in the video. The tracking data was analyzed by comparing the “ground truth” microbead position (computed from the image metadata) to the tracked microbead position (computed from the tracking algorithm). The tracking error magnitude was computed by computing the RMSE with all pairs of “ground truth” and tracked microbead positions.

Tissue tracking accuracy was evaluated by real-time tracking the displacement of tissue during an injection experiment. First, the user loaded a piece of human brain organoid tissue into the microscope, manually annotated the tissue edge for injection, and then proceeded with the injection trial. During the trial, the ML-guided Autoinjector tracked the tissue displacement and saved a camera image and the tracked tissue position after every time the pipette penetrated the tissue. After the injection experiment, the user manually re-annotated the tissue edge in each of the saved images. These manually re-annotations were used as the “ground truth” annotation positions for the displaced tissue. The tracking data was analyzed by comparing the “ground truth” tissue position (from the manual annotations) to the tracked tissue position (computed from the tracking algorithm). The tracking error magnitude was computed by computing the RMSE with all pairs of “ground truth” and tracked microbead positions.

Tissue tracking injection success was evaluated by performing a series of injection trials with different combinations of tracking on/off and manual/automatic annotations. In each injection trial, the total number of attempted injections was recorded. Injection success was assessed post-injection by imaging each tissue section on a confocal microscope and counting the number of successfully injected cells. Injection success percentage is the successful injections divided by attempted injections.

Tissue tracking injection duration was evaluated by performing numerous injection trials with tracking on/off. A video was recorded of each injection trial, and the duration was assessed post-injection by analyzing the start- and end-timepoints of the injection process. Trials without tracking were compared to trials with tracking.

---

**SUPPLEMENTARY FIGURE 1.**
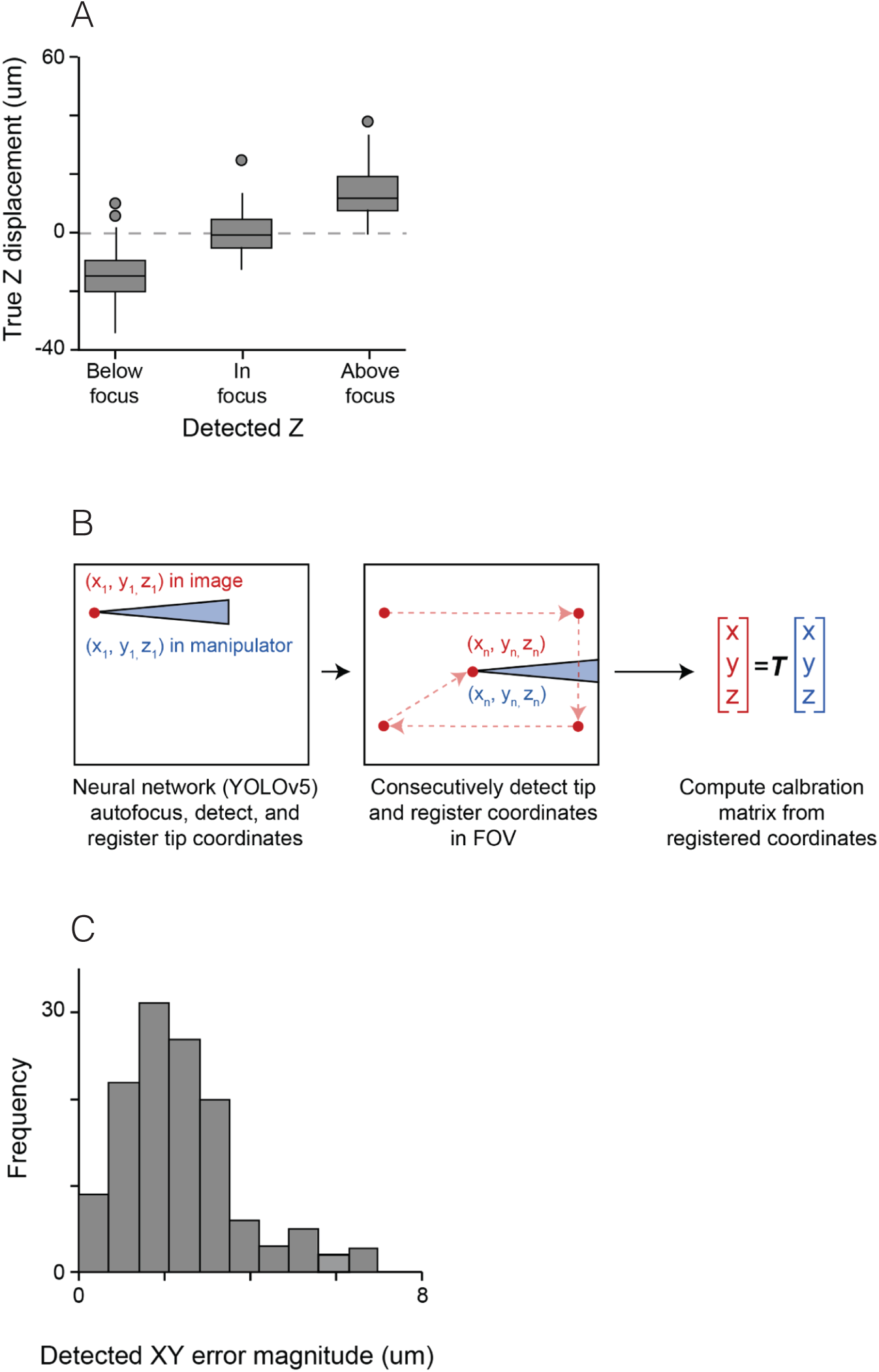
(A) Characterization of pipet time focus detection algorithm. (B) Schematic representation of ML object detection-based calibration procedure that allows movement in pixel coordinate space to be transformed to movement in the manipulator cartesian coordinates. (C) Graph representing the frequency of detected XY error magnitude after calibration

**SUPPLEMENTARY FIGURE 2.**
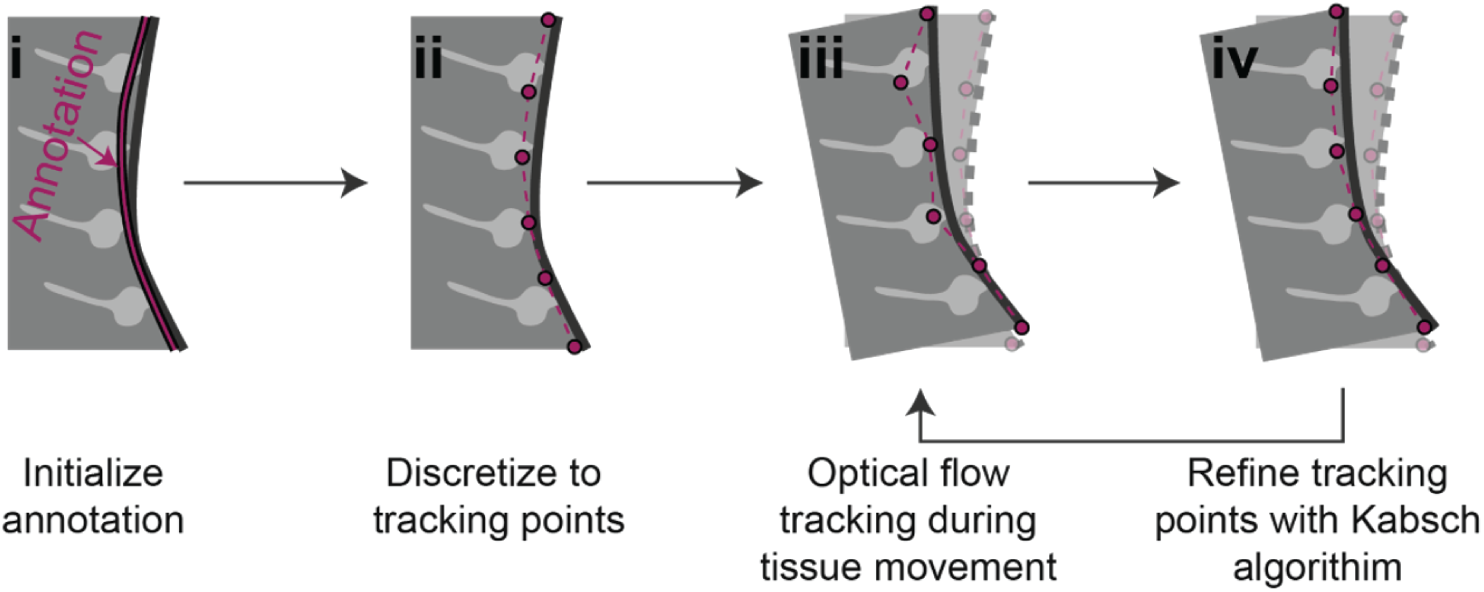
Schematic representation of the tissue displacement compensation algorithm implemented in the ML-guided Autoinjector robot.

## ACKNOWLEDGEMENTS

We are grateful to the services and facilities of HT for the outstanding support provided, notably, N. Maghelli and the team of the National Facility (NF) or Light Imaging, G. Faga’ and the team of Genome Editing and Disease Modeling and M. Ventura and M. Capillo and the team of Preclinical Research Facility. We thank G. Faga’ and P.Gatti and F. Martino (HT) for the critical reading of the manuscript and all members of the Taverna lab for helpful discussions.

## AUTHOR CONTRIBUTIONS

M.P. and J.O.B equally contributed to this work. M.P. worked on the experimental part and on organoids protocol set up. J.O.B wrote the code and worked on the robotic device set up. E.R. worked on and supported with the experimental part. M.P. is a Ph.D. student within the European School of Molecular Medicine (SEMM).

